# Looking Across Protein Domains to Identify Driver Mutations in Cancer

**DOI:** 10.1101/2025.08.11.669561

**Authors:** Daria Ostroverkhova, Yiru Sheng, Igor Rogozin, Anna R. Panchenko

## Abstract

Cancer can develop through the accumulation of somatic mutations that drive uncontrolled cell proliferation. A central objective in cancer research is to identify mutations that provide a selective growth advantage to tumor cells, so called driver mutations. Many computational methods infer driver missense mutations in proteins by assessing their recurrence. However, such approach suffers from the limited capacity to detect those driver mutations that occur infrequently across tumor samples. One strategy to overcome this limitation is to aggregate mutations from proteins sharing the same protein domain. Here we constructed a benchmark of cancer driver and passenger mutations, based on the known experimental and clinical studies, and systematically evaluated the applicability of methods that aggregate mutations across different mutation types and protein domains. We found that accounting for evidence mutations from different types of amino acid substitutions occurring in the same protein position enhances the classification performance. Furthermore, accounting for evidence mutations from paralogous proteins in the domain family increased the precision but compromised the overall classification accuracy. In addition, the performance of domain-based approaches was shown to crucially depend on the similarity between the target and evidence proteins.

## Introduction

Cancer may arise from the accumulation of somatic mutations that disrupt cellular homeostasis and promote uncontrolled proliferation (1). One of the central goals in cancer research is the identification of mutations that directly contribute to cancer progression. These are so-called driver mutations (2). A number of computational methods have been developed to distinguish driver mutations from passenger mutations, reviewed in (3). Many of these approaches use the recurrence of mutations within individual genes (a gene-based approach) across tumor samples as an indicator of driver status. However, this gene-based framework has limited capacity to detect rare driver mutations that occur infrequently across tumor samples. As a result, many low-frequency driver mutations might remain undetected due to a lack of statistical power even though most cancer driver mutations are rare (4,5). One strategy to overcome this limitation is to aggregate mutations from certain protein regions or protein domains (6-16).

The key idea underlying the domain approach is that driver mutations tend to affect aligned and equivalent amino acid positions across different proteins that share the same homologous domain family. Such equivalent positions are identified through the multiple sequence alignment (MSA) of proteins sharing the same domain family. By comparing the distribution of mutations in the selected region with that of the rest of the protein domain, such domain based approaches determine whether there is an enrichment in a number of mutations in specific MSA positions. Mutations affecting these positions may confer a selective advantage to cancer cells. In contrast, passenger mutations are expected to be randomly distributed and less likely to occur at the same position across the domain alignment. By aggregating mutations across domain family members, this approach may increase the statistical power potentially allowing the detection of low-frequency driver mutations. By doing so, the functional and cancerogenic consequences of uncharacterized mutations (whether they are rare or not) can be inferred based on the corresponding aligned positions harboring well-characterized driver mutations in other proteins that contain the same domain (6,7,17,18).

Domain-family based approach assumes that equivalent positions within a domain have similar functional and possibly cancerogenic effects. However, cancer type, mutant allele and amino acid type specificity in cancer mutational distributions has been noted previously (19). Indeed, different tissues and cancer types may promote certain oncogenic mutations. Moreover, various paralogous proteins from the same domain family can be differentially expressed in various tissues and cancer types. As a consequence, many driver mutations may exert oncogenic effects only when present in particular protein paralogs. Prediction of driver mutations by inference across protein domain family can be questionable in certain cases but its reliability has not been systematically evaluated previously due to the lack of the robust benchmarks.

Here we developed several mutational scoring models that leverage mutation type and domain-family mutation statistics, integrating the frequency of observed mutations, the background mutation rate, and other factors, to determine whether individual mutations are potential drivers. We constructed a benchmark set of cancer driver and passenger mutations based on known experimental and clinical studies, and for the first time systematically evaluated the applicability range of methods that aggregate mutational observation frequencies.

## Materials and Methods

### Datasets of Cancer Mutations

The pan-cancer somatic mutation data was retrieved from a previous study (20). Briefly, this dataset comprised whole exome sequencing (WES) data from 12,989 cancer patients, obtained through the Genomic Data Commons (GDC) from The Cancer Genome Atlas, Multiple Myeloma Research Foundation CoMMpass, Human Cancer Model Initiative, Count Me In (CMI), and the Clinical Proteomic Tumor Analysis Consortium (CPTAC) projects. As the result, the pan-cancer dataset contained 1,567,062 missense mutations.

### Protein Domain Identification and Construction of Multiple Sequence Alignments

Full-length protein sequences of human proteins were retrieved from the Consensus Coding Sequence (CCDS) database (version 24) (21), selecting only those with a “Public” ccds_status to ensure their high quality and validation. The CCDS database provides annotated protein sequences. We used the longest isoform among all isoforms produced by a single gene, resulting in a total of 19,085 protein sequences.

To identify protein domain regions, the Pfam database (version 37.0) of human protein domains was downloaded from the Pfam FTP service (https://ftp.ebi.ac.uk/pub/databases/Pfam/current_release/Pfam-A.hmm.gz) (22). HMMER’s hmmsearch tool (version 3.4, (23)) was applied to annotate the Pfam domain boundaries in the obtained protein sequences, with an E-value cutoff of ≤ 10^−8^. Domain regions of all human protein sequences were extracted and grouped together if they corresponded to the same Pfam domain family. These grouped sequences were then aligned using the MUSCLE (version 3.8.31) multiple sequence alignment (MSA) program (24). Next, we mapped missense mutations from the pan-cancer mutation dataset onto the domain multiple sequence alignments for further analysis. If a mutation was mapped to two different Pfam domains, the one with the lowest E-value was used for the analyses.

### Scoring Schemes for Ranking Mutations with respect to their Driver Status

We designed several scoring schemes: those based on a single protein (the score names start with “*S”*) and those incorporating evidence mutations from other proteins in the domain alignment (the score names start with “*D”*). A target mutation *q (t,l,k)* is specified by the type of mutation *t* and by the position *l* in the target protein *k*. A “mutation type” refers here to the substitution of one amino acid with another (e.g., Ala → Tyr). The simplest single protein score (*tS-score)* is calculated as a sum of the observed numbers of a target mutation *q* in different tumors. Next, two types of additional evidence were taken into account. The first type of evidence comes from aggregating all mutation types, not just the target mutation, at a given position. The corresponding *S-*score sums over all evidence mutations of different types (forming a set *T*) at position *l* in the target protein *k*:

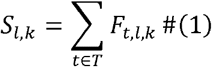

where *F*_*t,l,k*_ is the observed number of a mutation type *t* in a given position *l* in the target protein *k*. The second type of evidence comes from mutations observed in the aligned position *l* in other evidence proteins from the same domain family in the form of domain score:

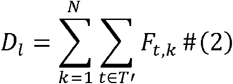

where *N* is the number of proteins containing at least one missense mutation at position *l. T’* is the set of all mutation types observed in the aligned column *l* of the MSA.

Further, we incorporated the background mutability of a mutated codon for a target protein:

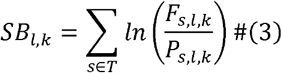

where *F*_*t,l,k*_ is the frequency of observed mutations of type *s* at a position *l* in the target protein *k. P*_*s,l,k*_ represents the codon mutability and is calculated as the probability of observing a specific type of codon change, defined by mutation type *s*, at a given position by chance, without selection. It was calculated according to formula (2) from (25).The corresponding domain score is calculated as:

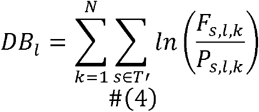

To penalize for proteins with no mutations at the aligned position, we normalized each score by dividing it by the number of non-gap characters in that position. We also tested scoring schemes including weighting by the similarity of mutation types between target and evidence mutations, but this did not improve the classification performance.

To compare the performance of the domain-based scores with the commonly used approach for identifying mutational hotspots within a domain family, we applied a binomial probability model as described in (7). The probability of observing *g* or more mutations at a specific position by chance, assuming mutations are uniformly distributed across a domain of length *L*, is computed using the binomial distribution:

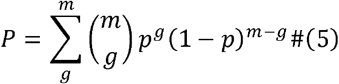

where 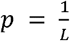 is the total number of mutations of any type in the domain family, and *m* is the total number of mutations in the domain.

### Construction of Benchmark Datasets

To evaluate the performance of different scoring models, we constructed a benchmark of annotated driver and passenger mutations by collecting the following data sets and combining them. The first dataset was obtained from a previous study (26) and contains experimental evidence of the impact of mutations, including changes in enzymatic activity, ligand binding response, downstream pathway effects, cellular transformation ability, tumor induction, and progression-free or overall survival in pre-clinical models. Mutations that did not affect normal protein function or tumor formation were classified as “passenger” mutations. From this benchmark, we retained only those mutations that were observed in the pan-cancer dataset and had available transcript identifier, yielding 588 labeled missense mutations: 494 drivers and 94 passengers.

We also utilized the MafAnnotator Python package provided by the OncoKB database (https://www.oncokb.org) to annotate mutations in the pan-cancer dataset. The OncoKB database is a precision oncology resource that provides manually curated annotations of the biological impact and clinical relevance of genetic variants in cancer (27,28). We filtered out those mutations which were labeled as “Inconclusive”, “Resistance”, and “Unknown”. The “Likely neutral” mutations were treated as passenger mutations, while “oncogenic” and “likely oncogenic” mutations were considered as drivers. As the result, annotated dataset contained 2,241 missense mutations: 2,016 driver and 225 passenger mutations.

Moreover, we used the Database of Cancer Passenger Mutations – the dbCPM database (4), a manually curated database cataloging experimentally validated cancer passenger mutations. For this study, we included only passenger mutations with “Level 1” annotations (validated by in vivo functional experiments) or “Level 2” annotations (validated by in vitro functional experiments) from the database. We then further included only those mutations that were observed in the pan-cancer dataset, resulting in 136 passenger mutations.

Furthermore, we merged these datasets to construct a combined benchmark. Driver and passenger labels for *POLE* and *POLD1* mutations were also obtained from a previous study (29). Any conflicting labels between datasets were removed. As a result, the final combined benchmark included 2,576 labeled missense mutations, comprising 2,037 drivers and 539 passengers. The final benchmark is available at https://github.com/Panchenko-Lab/Benchmark/blob/main/cancer_mutations_benchmark_updated.csv.

We then mapped the missense mutations from the benchmark onto the domain alignments, resulting in 27.1% (146) of passenger mutations and 45.8% (932) of driver mutations being successfully mapped to the alignments. After mapping mutations on protein domain families, we observed that some aligned columns (9.8%) exhibited inconsistent annotations: some proteins in the same column harbored both driver and passenger mutations, but most likely this value is underestimated, given the limited number of passenger mutations in our data set.

### Evaluation of Scoring Models Using Benchmark Datasets

To quantify the performance of different scoring schemes, we performed the ROC and precision-recall analyses. To address imbalances in the labeled datasets, we also calculated the maximal Matthew’s correlation coefficient (MCC). At the maximal value of MCC, we derived the confusion matrix and calculated overall accuracy, precision, specificity, sensitivity, and F1 score. Furthermore, the error rate (ERR) was computed at maximal MCC as ERR = (FP+FN)/(TP+FP+TN+FN). Here FP refers to false positives, namely, mutations predicted as drivers but labeled as passengers in the benchmark dataset, whereas FN denotes false negatives, referring to true driver mutations that were incorrectly predicted as passengers. TP represents true positives, i.e., driver mutations correctly predicted as drivers, and TN correspond to true negatives, i.e., passenger mutations correctly predicted as passengers.

As described above, missense mutations were first mapped onto all proteins from the MSAs of protein domain families. Each mutation with the benchmark label (“driver” or “passenger”) was treated as a target mutation, while the remaining mutations served as evidence mutations for that target. We then calculated all scores for each target mutations. To estimate the error in evaluation metrics, we applied the bootstrapping process to this list of target mutations. We randomly sampled, with replacement, an equal number of driver and passenger mutations from the list of labeled target mutations to construct each bootstrap sample. This approach ensured that each sample was class-balanced, despite the overall labeled list being imbalanced toward driver mutations. A total of 100 bootstrap samples were generated. For each bootstrap sample, evaluation measures were then computed, and the resulting values across all samples were used to calculate mean values and corresponding 95% confidence intervals.

Next, we evaluated the performance of scoring models across different ranges of sequence identity between target and evidence sequences. Because of the limited number of benchmark mutations, we assessed the performance of each scoring scheme at two sequence identity ranges — below 50% and above 50% — using pairwise comparisons between the protein carrying the target mutation and each protein carrying evidence mutation in the MSA.

## Results and Discussion

### Variable Impact of Additional Mutational Evidence on Classification Accuracy

We have evaluated and compared the performance of different scoring models. An overview of the study framework is shown in Figure 1. A total of 92 domain families, each containing mutations from more than one evidence protein and at least one mutation with a benchmark annotation, was included in the analysis. First, we evaluated the overall classification accuracy in terms of distinguishing drivers from passenger and found that aggregating mutations of all types, rather than only the target mutation type, considerably improved the classification accuracy (MCC = 0.55 and 0.37 for S-score and tS-score respectively, Table 1). Second, the domain-based scores designed in this study achieved higher performance, compared to mutational hotspot identification methods that use Binomial statistics, in the low range of sequence similarity between target and evidence proteins (Figure 3C). A similar trend was noted when we evaluated the scoring models on rare mutations that were observed only once in cancer patients (Supplementary Table 2). However, for rare mutations aggregating across different mutation types in a single protein was beneficial for predicting passenger mutations (specificity of 0.96) to the larger degree compared to predicting driver mutations (sensitivity of 0.44). In terms of precision, defined as a proportion of predicted drivers that are truly labeled as drivers, the domain-based scores yielded higher precision (0.96) than single protein scores (0.86), indicating that the domain scoring scheme correctly identifies the most prominent driver mutations. Indeed, as shown in Figure 2A-B, there were two prominent mutational hotspots in the Furin-like family and four mutational hotspots in the RAS family. Five of these hotspots had supporting evidence mutations from other proteins at the same aligned position.

**Table 1.** Performance of different mutation scoring strategies in identifying driver mutations with the corresponding confidence intervals calculated by bootstrapping.

| **Score Name** | **Max MCC** | **ROC AUC** | **Precision at Max MCC** |
| --- | --- | --- | --- |
| **tS** | $0.356 \pm 0.005$ | $0.693 \pm 0.003$ | $0.833 \pm 0.008$ |
| **S** | $0.554 \pm 0.008$ | $0.828 \pm 0.004$ | $0.84 \pm 0.007$ |
| **SB** | $0.587 \pm 0.008$ | $0.819 \pm 0.005$ | $0.859 \pm 0.007$ |
| **D** | $0.366 \pm 0.009$ | $0.678 \pm 0.006$ | $0.852 \pm 0.017$ |
| **DB** | $0.319 \pm 0.007$ | $0.609 \pm 0.007$ | $0.961 \pm 0.014$ |
| **Binomial** | $0.262 \pm 0.011$ | $0.634 \pm 0.007$ | $0.628 \pm 0.011$ |

Notes: tS, type-specific single protein score; S, single protein score; SB, single protein score with background mutability; D, domain-family score; DB, domain-family score with background mutability; Binomial, hotspot score based on binomial probability model; MCC, Matthew's correlation coefficient; ROC AUC, area under the receiver operating characteristic curve.

**Figure 1.**
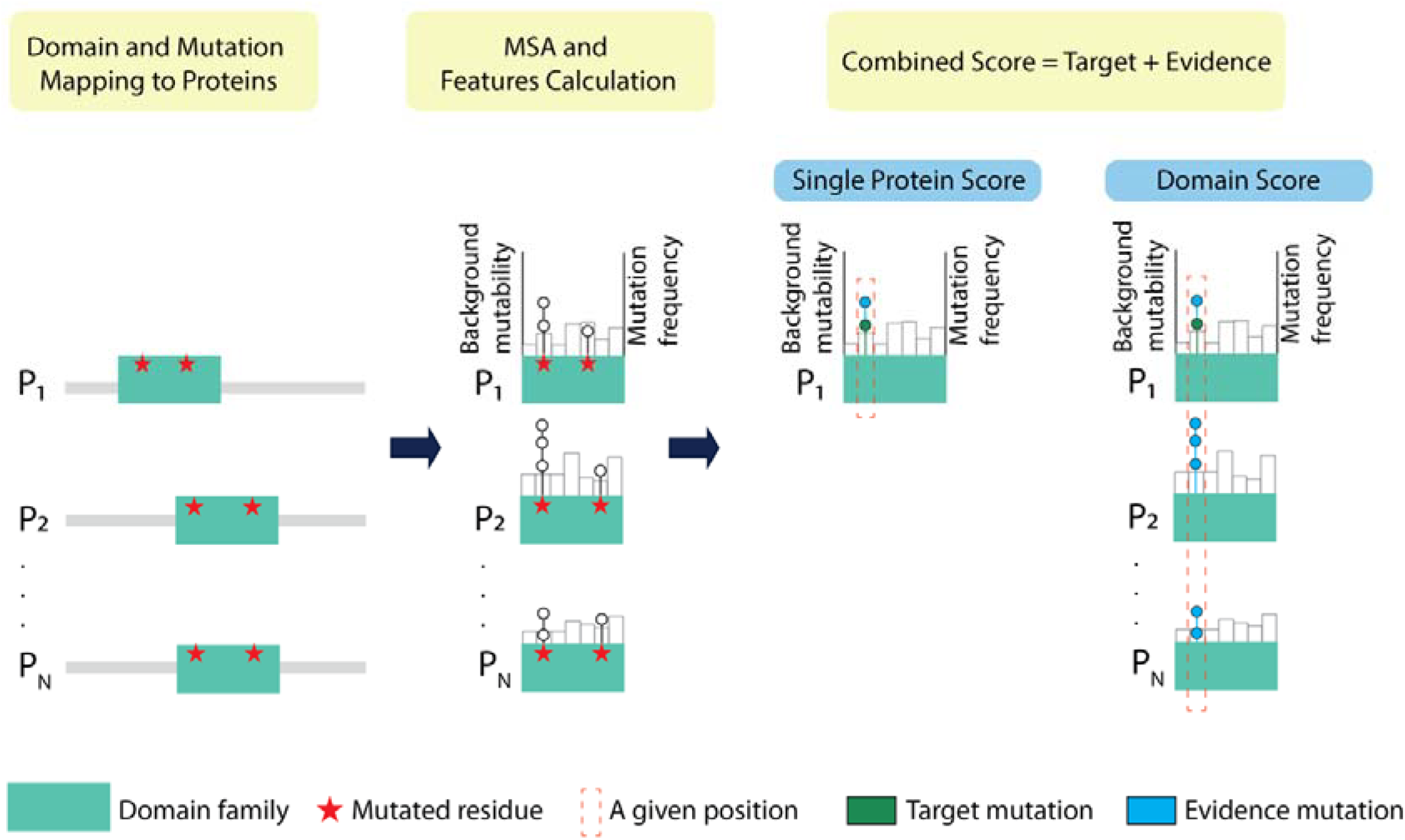
Schematic overview of the scoring framework. Domain annotations are first assigned to protein sequences, and missense mutations are then mapped to the corresponding domain regions within each protein. These mutations are subsequently mapped to the multiple sequence alignment (MSA) of the domain family, allowing background mutability (shown as empty boxes) and mutation frequency to be computed for each specific amino acid substitution at each aligned position on each protein. In each aligned column, the *target mutation* refers to a missense mutation of interest, and all other mutations at the same aligned position, whether from the same or other proteins, are considered *evidence mutations*. A mutation score is then computed for the target mutation by incorporating both its own features and those of the evidence mutations. In the visualization, each lollipop represents a missense mutation; its height reflects mutation frequency. The border color of each lollipop indicates whether it is a target (green) or evidence (blue) mutation.

**Figure 2.**
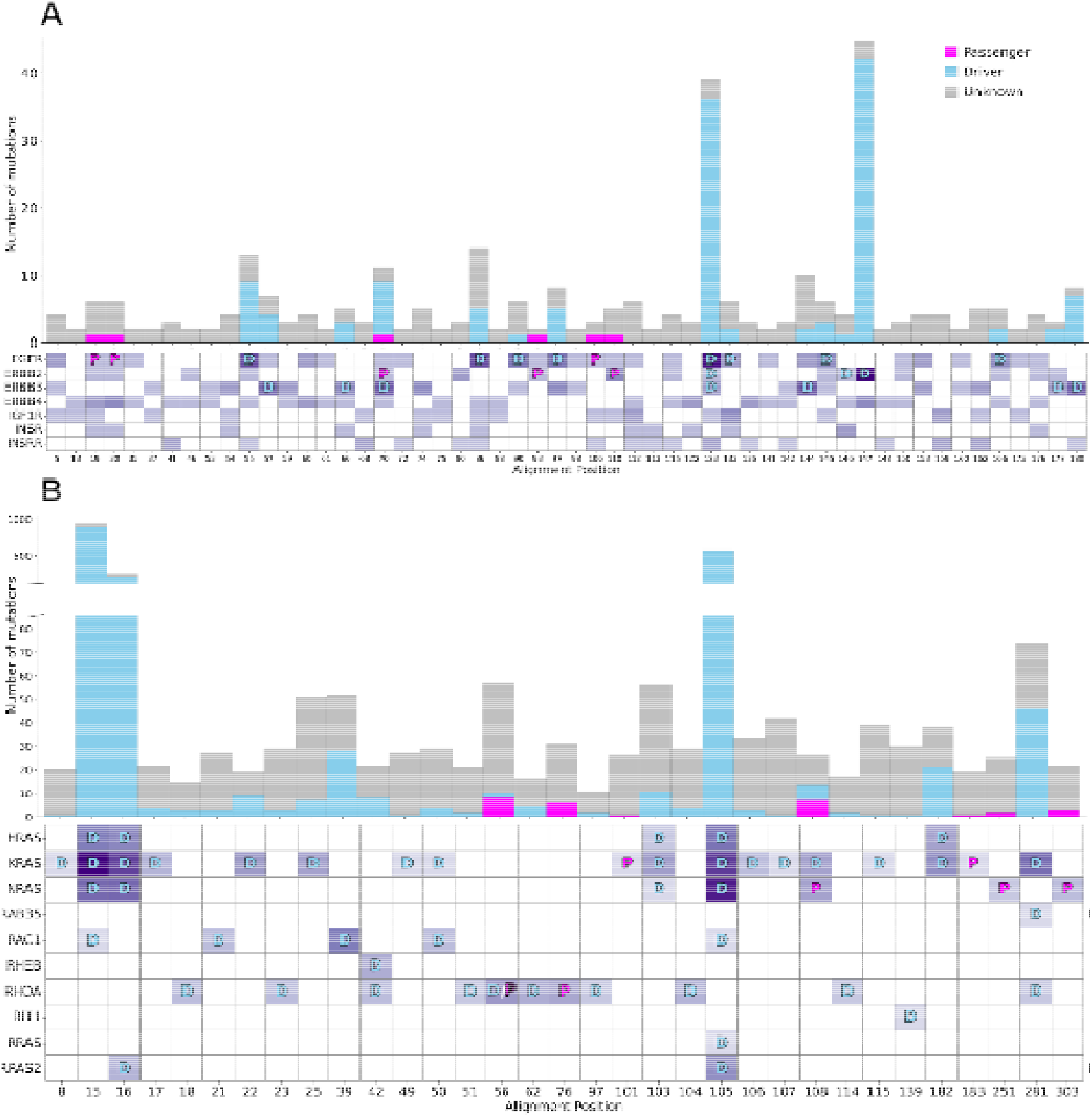
Mapping of mutations on the Furin-like and RAS domain families. **(A)** The stacked bar plot (top) shows the number of mutations per aligned position across members of the Furin-like domain. Only positions with at least one mutation are shown. Bars are colored according to the benchmark mutation label: Driver, Passenger, or Unknown. The heatmap (bottom) shows the number of mutations across proteins in the domain family and the color intensity reflects the number of mutations observed at each position. Each row represents a protein, and each column corresponds to an aligned position in the multiple sequence alignment. Cells labeled **D** indicate driver mutations and **P** indicate passenger mutations. Only those columns containing at least one labeled mutations are shown. **(B)** The same is shown for RAS domain family. The complete heatmap of the RAS domain is provided in Supplementary Figure 1.

**Figure 3.**
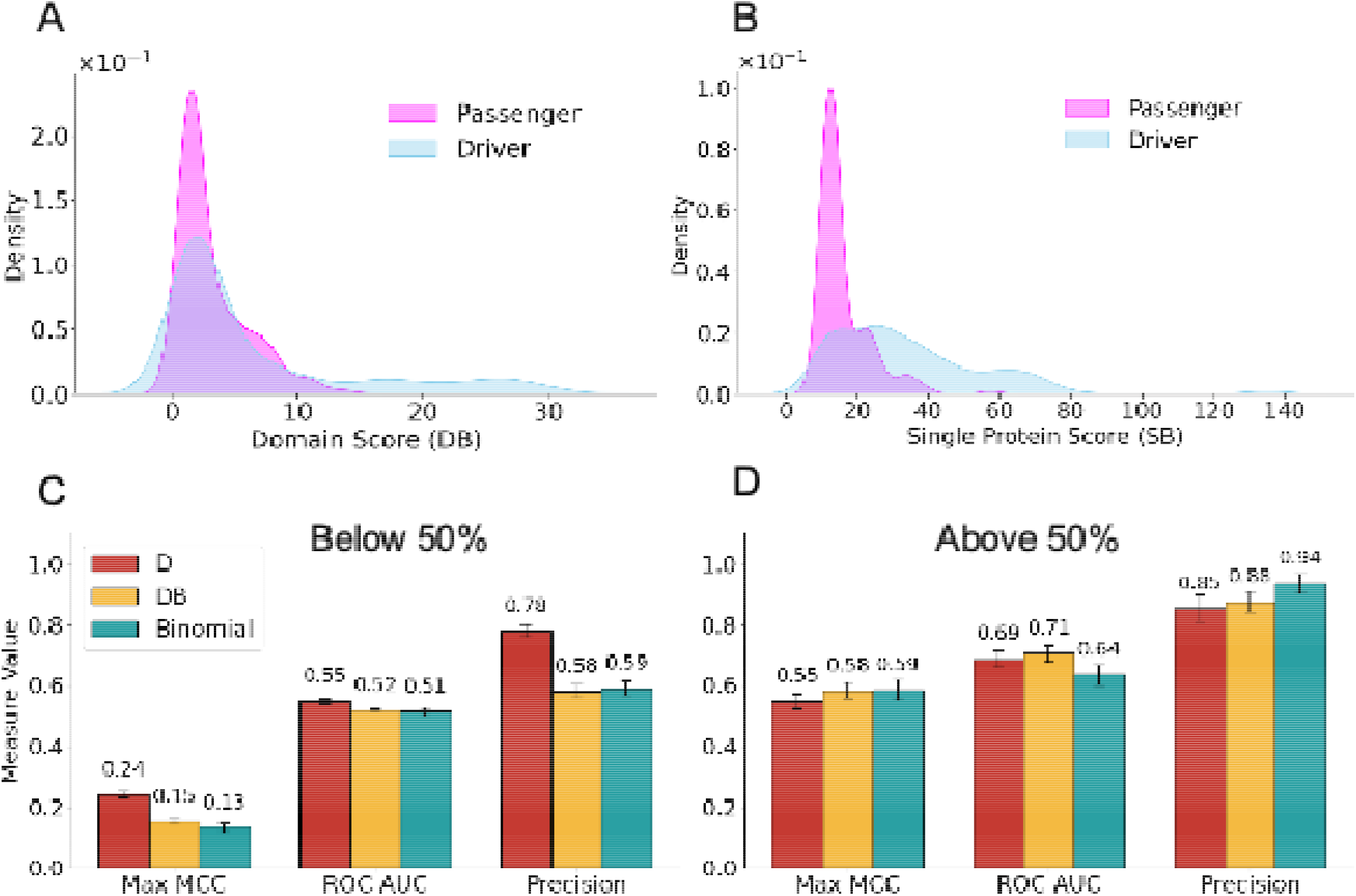
Performance evaluation of domain and single protein based scoring schemes for driver mutation identification. The distribution of Domain Score (DB) **(A)** and Single Protein Score (SB) **(B)** for driver and passenger mutations. Performance comparison of three scoring schemes (D, DB, and Binomial statistics) evaluated on benchmark data for two sequence identity ranges between target and evidence proteins: 1–50% **(C)** and 50–100% **(D)**. Bar plots show Maximal Matthews Correlation Coefficient (Max MCC), Area Under the ROC Curve (ROC AUC), and Precision, with error bars indicating 95% confidence intervals. D, domain-family score; DB, domain-family score with background mutability; Binomial, hotspot score based on binomial probability model.

However, other than mutational hotspots, no matter which scoring models we used, incorporating mutational data from a protein domain did not boost the overall classification performance, even when accounting for different mutation types or background mutability. Namely, the maximal classification performance was MCC = 0.37 for domain score, whereas the single-protein scoring scheme yielded a higher MCC value of 0.59. Indeed, as shown in Figure 3A-B, the distributions of domain scores for passengers and drivers substantially overlap, unlike the ones of single protein scores. Possible reasons of such limited accuracy in case of domain scores are illustrated in Figure 2. Namely, positions harboring passenger mutations can be inflated by unannotated evidence mutations (in grey) making passenger mutation scores comparable to those of drivers. Some aligned positions contain both driver and passenger annotations. In addition, cancer driver mutations are not uniformly distributed among protein sequences in the domain alignment. Even though there are several prominent mutational hotspots, the vast majority of mutations are protein-specific, especially rare ones.

In addition, we conducted a separate analysis for each domain family, to assess whether incorporating evidence mutations from other proteins within the same family affects the classification performance. Out of 92 protein domain families, eight families that contained at least five mutations of each class (driver and passenger) were included in this analysis. The results were consistent with the overall trend of lower performance for domain-based scores compared to single-protein scores (Supplementary Table 3), with a few exceptions. For example, domain-based score in the RAS family achieved a slightly higher MCC (0.74) compared to scores based on a single protein (0.71).

### Similarity between the Target and Evidence Proteins is a Crucial Factor

It is important to assess if models’ performance is affected by the sequence identity between the target and evidence proteins. Our analysis showed that prediction accuracy was strongly influenced by the sequence identity to the target protein. In particular, the MCC value for the domain score was 0.58 for 50-100% identity range (Figure 3D), compared 0.24 for 0-50% identity range (Figure 3C-D, Supplementary Table 4-5). Indeed, higher sequence similarity between proteins tends to correspond to greater similarity in their specific biological functions. However, the relationship between the shared sequence and structural features of protein domains and their function is complex – particularly in evolutionarily and functionally diverse families (30-32). This complexity is further supported by findings from other systems, which demonstrate that mutations at equivalent residues can yield divergent phenotypic and disease outcomes (33).

### Conclusion

Overall, our findings indicate that leveraging additional evidence mutations, whether from different substitution types in the same protein or from different homologous proteins, can variably impact classification performance of drivers versus passengers, and depends on the specific task. For applications focused on identifying a limited number of high-confidence driver mutations with high accuracy, incorporating evidence mutations is advantageous. However, this strategy may be less effective for detecting unknown or rare drivers. We show that the key determinants of performance might include protein specificity and the degree of similarity between the target and evidence proteins. It is also important to note that mutations affecting mutational hotspots tend to be more prevalent in oncogenes, whereas tumor suppressor genes often exhibit more dispersed mutational patterns. We also acknowledge the limitations of our analysis. While this study offers the thorough evaluation of driver and passenger mutation identification and its modulation by supplementary evidence, it is nonetheless limited by the incompleteness of current driver and passenger annotations and limited family representation. Largest families of cancer related paralogous proteins, such as histones, could not be studied, exactly because of the lack of such annotations.

## Acknowledgements

Regretfully, one of the co-authors, Igor Rogozin, passed away tragically and far too soon while this paper was being prepared for submission. The authors, and especially ARP, wish to express their deep gratitude for Igor’s invaluable contributions to discussions and his innovative ideas. He will be greatly missed by his colleagues.

The authors were supported by the Department of Pathology and Molecular Medicine, Queen’s University, Canada. A.R.P. is the recipient of a Senior Canada Research Chair in Computational Biology and Biophysics. This study was conducted with the support of the Ontario Institute for Cancer Research through funding provided by the Government of Ontario. I.B.R. was partially funded by the EU’s Operational Program “Just Transition” CZ.10.03.01/00/22_003/0000003 LERCO.

