## Supplementary Figure 1 for "Looking Across Protein Domains to Identify Driver Mutations in Cancer"

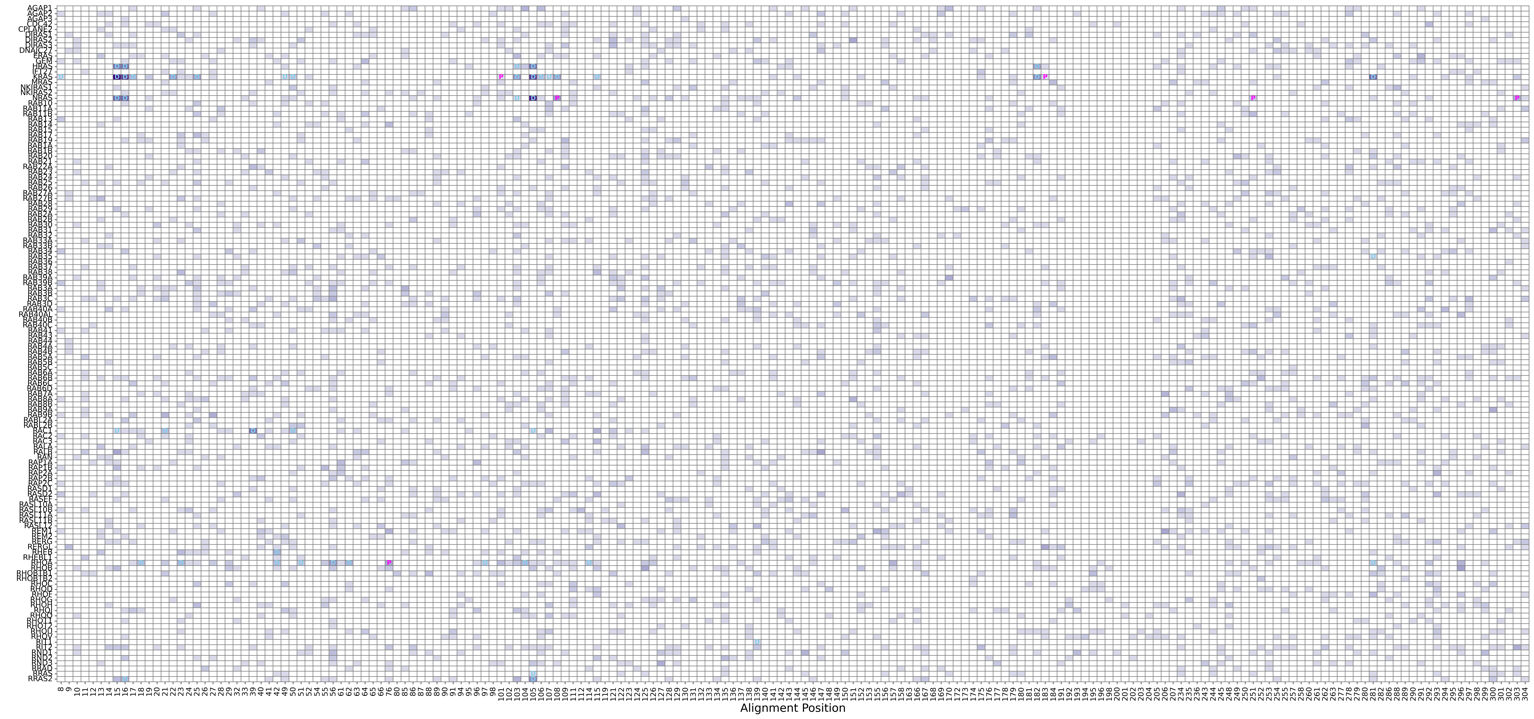

**Supplementary Figure 1.** The heatmap of mutations across all proteins in the RAS domain family. Each row represents a protein, and each column corresponds to an aligned position in the multiple sequence alignment. Mutations are labeled as driver (D) or passenger (P), and the color intensity reflects the number of mutations observed at each position.
